# Bias, robustness and scalability in differential expression analysis of single-cell RNA-seq data

**DOI:** 10.1101/143289

**Authors:** Charlotte Soneson, Mark D. Robinson

## Abstract

**Background:** As single-cell RNA-seq (scRNA-seq) is becoming increasingly common, the amount of publicly available data grows rapidly, generating a useful resource for computational method development and extension of published results. Although processed data matrices are typically made available in public repositories, the procedure to obtain these varies widely between data sets, which may complicate reuse and cross-data set comparison. Moreover, while many statistical methods for performing differential expression analysis of scRNA-seq data are becoming available, their relative merits and the performance compared to methods developed for bulk RNA-seq data are not sufficiently well understood.

**Results:** We present *conquer*, a collection of consistently processed, analysis-ready public single-cell RNA-seq data sets. Each data set has count and transcripts per million (TPM) estimates for genes and transcripts, as well as quality control and exploratory analysis reports. We use a subset of the data sets available in *conquer* to perform an extensive evaluation of the performance and characteristics of statistical methods for differential gene expression analysis, evaluating a total of 30 statistical approaches on both experimental and simulated scRNA-seq data.

**Conclusions:** Considerable differences are found between the methods in terms of the number and characteristics of the genes that are called differentially expressed. Pre-filtering of lowly expressed genes can have important effects on the results, particularly for some of the methods originally developed for analysis of bulk RNA-seq data. Generally, however, methods developed for bulk RNA-seq analysis do not perform notably worse than those developed specifically for scRNA-seq.

## Background

High throughput sequencing of cDNA, or RNA-seq, has been used routinely for transcriptomic characterization and analysis for almost a decade [1, 2, 3], but until recently, technical limitations have confined researchers to sequencing libraries prepared from pools of thousands or more cells. This restricts the analyses to *average* abundance levels across all the cells in the library, and thus prohibits studies of transcriptome-wide heterogeneity of cell populations. However, recent technological advances now allow researchers to prepare sequencing libraries from minute amounts of RNA, and thus to profile the transcriptomes of individual cells [4, 5, 6, 7, 8]. Termed *single-cell RNA-seq* (scRNA-seq), this holds great promise in many subdisciplines of biology and medicine such as oncology, immunology and early development. With the adoption of scRNA-seq for more and more studies, the number of data sets deposited in public repositories such as the Short Read Archive (SRA, https://www.ncbi.nlm.nih.gov/sra)[9], Gene Expression Omnibus (GEO, https://www.ncbi.nlm.nih.gov/geo)[10, 11] and the European Nucleotide Archive (ENA, http://www.ebi.ac.uk/ena)[12], increases rapidly. The publicly available data is a highly useful resource for the research community, both for computational method development and for reanalysis and extension of published observations. For most public data records, both the raw read files and a processed data table, typically containing estimated gene abundances used by the data generators for their analyses, are available. Since the aims of different studies vary widely, naturally, different public data sets have often been processed using very different pipelines. Furthermore, the processed abundances may be represented in different units, such as read counts, transcripts per million (TPM) or fragments per kilobase per million mapped reads (FPKM), and sometimes a fraction of the cells and/or genes are filtered out due to poor quality or low abundance. Taken together, this can make reuse of the preprocessed public data sets, and especially comparisons across data sets, challenging. To simplify this aspect, we developed *conquer*, a collection of consistently processed, analysis-ready public scRNA-seq data sets collected using full-length transcript library preparation protocols. For each data set, we estimate abundances using Salmon [13], and provide both TPMs and estimated counts for each annotated gene and transcript. In addition, *conquer* provides reports summarizing the quality assessment and exploratory analysis of each data set to help users determine whether (a subset of) a particular data set is suitable for their purposes.

Arguably the most commonly performed computational task for RNA-seq data to date is differential gene expression analysis, that is, searching for genes showing differences in abundance associated with a given phenotype. While several well-established tools exist for differential expression analysis of bulk RNA-seq data (e.g., [14, 15, 16]), corresponding methods for scRNA-seq data are just emerging, and the relative merits of each of the existing methods, as well as their advantages compared to methods developed for bulk RNA-seq analysis, have not been fully investigated. Due to the special characteristics of scRNA-seq data, including generally low library sizes, high noise levels and a large fraction of so-called “dropout” events, that is, failure to observe some genes although they are indeed expressed in a given cell, it is unclear whether differential expression methods developed for bulk RNA-seq data, lacking these characteristics, are applicable also to scRNA-seq data. A few studies, starting to evaluate these aspects, have been published recently [17, 18]. Jaakkola *et al*. [17] compared five differential expression methods and concluded that neither the methods developed for bulk RNA-seq nor those specifically developed for scRNA-seq analysis performed well, but that their more general method (ROTS), originally developed for microarray data, showed good performance throughout their comparisons. Miao and Zhang [18] compared 14 differential expression methods, four of which were developed specifically for scRNA-seq data, and illustrated that the optimal method choice depends on the number of cells in the data set and the strength of the expected signal. In this study, we use several of the processed data sets available in *conquer* to empirically evaluate and compare differential expression methods, including both methods developed for bulk RNA-seq and methods developed explicitly for scRNA-seq data. Our study extends the previous comparisons to a larger set of scRNA-seq differential expression methods and a larger number of experimental data sets with different characteristics, and additionally includes evaluations based on simulated data. Moreover, we run several of the methods with different settings, investigate the effect of filtering out lowly expressed genes and extend the set of employed evaluation criteria. We focus on the simplest situation, contrasting two groups of cells and considering all cells within a group to be independent replicates, since this setup can be accommodated by all compared methods. However, it should be noted that some scRNA-seq data sets will contain cells from multiple biological replicates, or from multiple plates, introducing a hierarchical variance structure that is not accounted for by such a simple model [19]. Moreover, having data with single-cell resolution allows the researcher to ask additional questions that can not be addressed with bulk RNA-seq data, such as explicitly testing whether the different groups of cells show different levels of variability or multimodality [20, 21].

## Methods

### conquer

The *conquer* pipeline processes (sc)RNA-seq data sets obtained with full-length transcript protocols using the following steps:

1. Build a quasi-mapping transcriptome index for Salmon [13] from the combined set of annotated cDNA and ncRNA sequences as well as ERCC spike-in sequences. The Ensembl catalog (v38) [22] was used to process the currently available data sets.
2. For each scRNA-seq sample (cell):

(a) Find the corresponding SRA run ID(s) and download and merge the respective FASTQ file(s). If required, trim adapters using cutadapt [23].
(b) Perform quality control of the FASTQ file(s) using FastQC (http://www.bioinformatics.babraham.ac.uk/projects/fastqc/).
(c) Estimate transcript abundances (TPMs and estimated counts) using Salmon.
3. Summarize FastQC and Salmon diagnostics for all samples in the data set using MultiQC [24].
4. Read transcript abundances with the tximport R package [25] and create a MultiAssayExperiment object [26] containing gene- and transcript-level estimated counts and TPMs for the entire data set, as well as phenotypic information obtained from the public repository. For each gene, we include both aggregated transcript counts and length-scaled TPMs [25], as calculated by tximport.
5. Perform quality control, exploratory analysis and visualization of the gene-level abundances using the scater Bioconductor package [27] and summarize in a report.

Many of the processed data sets contain not only scRNA-seq samples, but also bulk RNA-seq samples for comparison, or technical control samples. Whenever these could be identified, they are excluded from the processed data. A list of the excluded samples is provided in the online repository. Cells belonging to the same SRA/GEO data set but sequenced on different platforms are separated into different data sets. No filtering based on poor quality or low abundance is performed, since that may introduce unwanted biases for certain downstream analyses and since no universally adopted filtering approach or threshold currently exists. However, the provided quality control and exploratory analysis reports can be used to determine whether some samples need to be excluded for specific applications. Since TPMs and counts are estimated using the same reference annotation, with the same software and using the same data, the *conquer* data sets can be used to compare computational methods that require different types of input, with minimal bias. The processed data sets and the resulting reports can be browsed and downloaded from http://imlspenticton.uzh.ch:3838/conquer/, and the underlying code used to process all data sets is available from https://github.com/markrobinsonuzh/conquer.

## Evaluation of differential expression methods

### Experimental and simulated data

Six of the real data sets from *conquer*, with a large number of cells, are selected as the basis for the evaluation of differential expression analysis methods. For each of the data sets, we retain only cells from two of the annotated cell types (Table 1), attempting to select large and relatively homogeneous populations among the ones annotated by the data generators. The selected data sets span a wide spectrum of signal strengths and population homogeneities (Supplementary Figures 1 and 2). For each data set, we then generate one instance of “maximal” size (with the number of cells per group equal to the size of the smallest of the two selected cell populations) and several subsets with fewer cells per group by random subsampling from the maximal size subset (see Table 1 for exact group sizes). For each non-maximal sample size, we generate five replicate data set instances, and thus each original data set contribute 11 or 16 separate instances, depending on the number of different sample sizes (Table 1). Moreover, for each data set with enough cells we generate *null* data sets with different sample sizes (again, five instances per sample size except for the maximal size) by sampling randomly from one of the two selected cell populations. Finally, three of the data sets (GSE45719, GSE74596 and GSE60749-GPL13112) are used as the basis for simulation of data using a slightly modified version of the *powsim* R package [28]. Individual reports verifying the similarity between the simulated and real data sets across a range of aspects are provided as supplementary material. As for the original, experimental data sets, we subsample data set instances with varying number of cells per group, and further generate null data sets by random sampling from one of the simulated groups. In each simulated data set, 10% of the genes are selected to be differentially expressed between the two groups, with fold changes sampled from a Gamma distribution with shape 4 and rate 2. The direction of the differential expression is randomly determined for each gene, with equal probability of up- and downregulation. Mean and dispersion parameters used as basis for the simulations are estimated from the respective real data sets using edgeR [15]. For each of the three data sets, the rounded length-scaled TPMs for all genes with at least two non-zero counts are used as input to the simulator, and a data set with the same number of genes is generated. The counts for each simulated gene are based on one of the original genes (however, the same original gene can be the basis for more than one simulated gene), and by retaining this information we can link average transcript lengths (calculated by tximport for the original data) to each simulated gene, and thus estimate approximate TPMs also for the simulated data.

**Table 1:** Experimental single-cell data sets used for the differential expression method evaluation.

| Data set | Compared cell subsets | Number of cells/group | Used for simulation | Organism | Ref |
| --- | --- | --- | --- | --- | --- |
| GSE45719 | 16-cell stage blastomere *vs* Mid blastocyst cell (92-94h post-fertilization) | 50, 24, 12, 6 | yes | mouse | [31] |
| GSE45719null | 16-cell stage blastomere | 24, 12, 6 |  | mouse | [31] |
| GSE48968-GPL13112 | BMDC (1h LPS Stimulation) *vs* BMDC (4h LPS Stimulation) | 95, 48, 24, 12 |  | mouse | [32] |
| GSE48968-GPL13112null | BMDC (1h LPS Stimulation) | 48, 24, 12, 6 |  | mouse | [32] |
| GSE60749-GPL13112 | v6.5 mouse embryonic stem cells, culture conditions: 2i+LIF *vs* v6.5 mouse embryonic stem cells, culture conditions: serum+LIF | 90, 48, 24, 12 | yes | mouse | [33] |
| GSE60749-GPL13112null | v6.5 mouse embryonic stem cells, culture conditions: 2i+LIF | 47, 24, 12, 6 |  | mouse | [33] |
| GSE74596 | NKT0 *vs* NKT17 | 44, 22, 12, 6 | yes | mouse | [34] |
| GSE74596null | NKT0 | 22, 12, 6 |  | mouse | [34] |
| EMTAB2805 | G1 *vs* G2M | 96, 48, 24, 12 |  | mouse | [35] |
| EMTAB2805null | G2M | 48, 24, 12, 6 |  | mouse | [35] |
| UsoskinGSE59739 | RT-1.NP1 *vs* RT-1.TH | 58, 36, 24, 12 |  | mouse | [29] |
| UsoskinGSE59739null | RT-1.NP1 | 24, 12, 6 |  | mouse | [29] |
| GSE63818-GPL16791 | Primordial Germ Cells, developmental stage: 7 week gestation *vs* Somatic Cells, developmental stage: 7 week gestation | 26, 12, 6 |  | human | [36] |

In addition to the six data sets from *conquer*, we downloaded and processed the UMI counts corresponding to the GEO entry GSE59739 [29] from http://linnarssonlab.org/drg/ (accessed December 18, 2016). The provided UMI RPMs are used in the place of TPMs, and were combined with the provided information about the total number of reads per cell to generate gene read counts. Empty wells were filtered out. For comparison, we also downloaded a bulk RNA-seq data set from the Geuvadis project [30] from http://www.ebi.ac.uk/arrayexpress/experiments/E-GEUV-1/ and estimated gene expression levels using the same pipeline as for the *conquer* data sets. For this data set, we perform differential expression analysis using a subset of the methods applied to the single-cell RNAseq data sets, comparing samples from the CEU and YRI populations generated at the University of Geneva.

For each real and simulated data set, we perform the differential expression analysis evaluation both on the full data set (excluding only genes with 0 counts in all considered cells) and on a filtered data set, where we retain only genes with an estimated TPM above 1 in more than 25% of the considered cells. Depending on the data set and the number of considered cells, between 20 and 50% of the genes are retained after this filtering (Supplementary Figure 3).

### Differential expression analysis methods

For each of the real and simulated scRNA-seq data sets, we apply 30 statistical approaches for differential expression analysis to compare the expression levels in the two groups of cells (Table 2). As representatives for methods developed for differential analysis of bulk RNA-seq data, we include edgeR [15], DESeq2 [14], voom-limma [16] and limma-trend [16]. For edgeR, we apply both the likelihood ratio test (LRT) [37] and the more recent quasilikelihood approach (QLF) [38]. For the LRT, in addition, we use both the default dispersion estimates [39] and the robust dispersion estimates developed to address outlier counts [40], and we apply edgeR both with the default TMM normalization [41] and with the recently developed deconvolution normalization approach for scRNA-seq [42]. DESeq2 is run in two modes, after rounding the length-scaled TPM values to integers: either with default settings, or disabling the internal independent filtering and outlier detection and replacement. Additionally, both edgeR/LRT and DESeq2 are applied to both the read counts (length-scaled TPMs as described above) and the recently suggested census transcript counts [43], which are calculated from the estimated TPMs using monocle [44] (v2.2.0) with default settings.

**Table 2:**
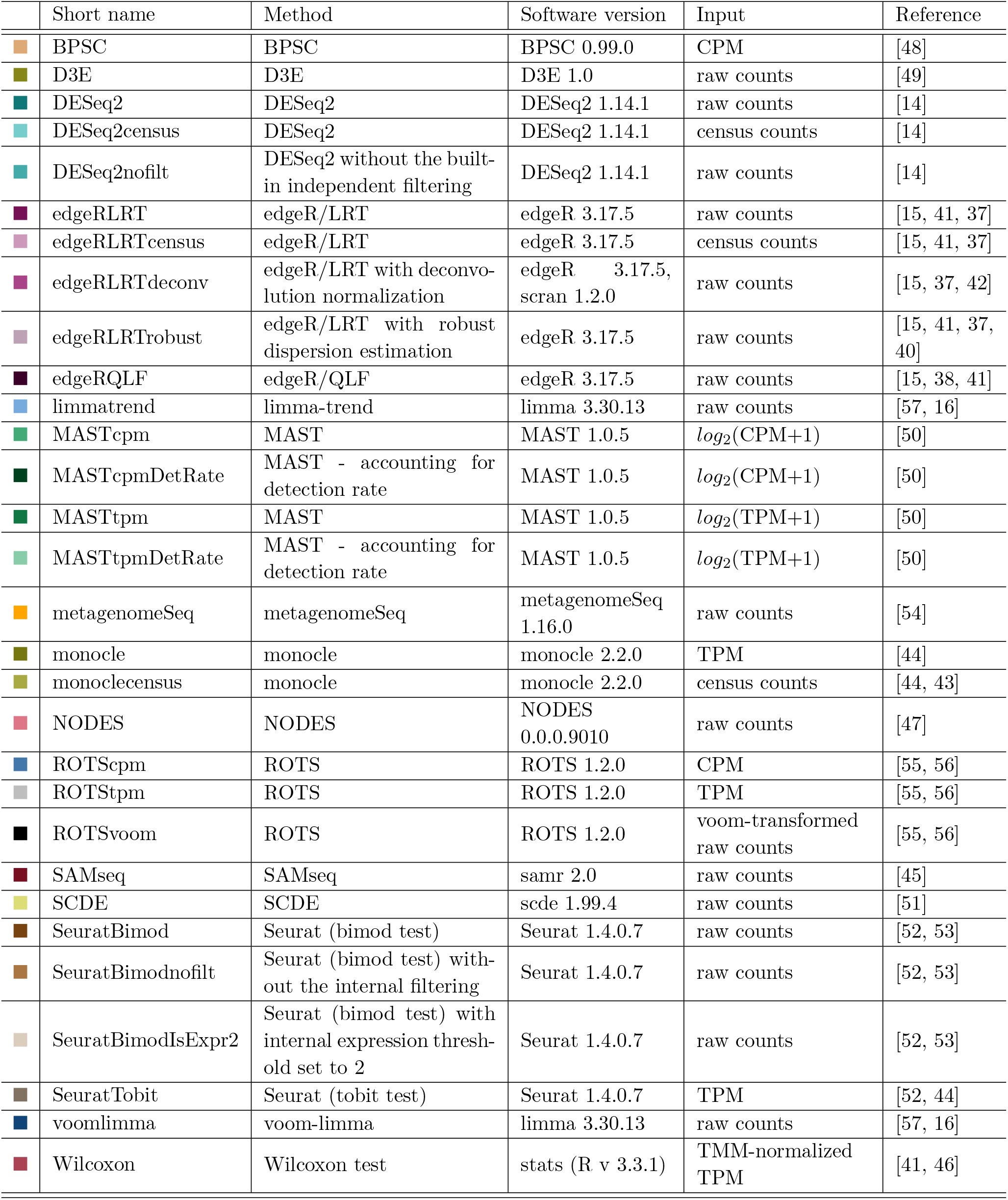
Evaluated differential expression methods, together with package versions and the type of input values provided to each of them. Note that “raw counts” here refers to length-scaled TPMs, which are on the scale of the raw counts, but are unaffected by differential isoform usage [25]. CPM values are calculated with edgeR (v 3.17.5), and census counts with monocle (v 2.2.0).

Three non-parametric methods are included in the comparison: SAMseq [45], the Wilcoxon test [46] and NODES [47]. SAMseq is applied to the length-scaled TPMs, while the Wilcoxon test is applied to TPM estimates after applying TMM normalization to address the compositionality of the TPMs. NODES was initially run in two modes: with default settings, and after disabling the internal filtering steps. However, disabling the internal filtering caused the method to fail in subsequent steps, and thus we retain only the runs with default settings.

We include a broad range of methods developed specifically for scRNA-seq differential expression analysis. BPSC [48] is applied to CPMs (calculated using edgeR) as suggested by the package authors. D3E [49] is run with the method-of-moments approach to parameter estimation, the non-parametric Cramer-von Mises test to compare distributions and without removing zeros before the analysis. MAST [50] is applied to both log_2_(CPM+1) and log_2_ (TPM+1) values, both with and without including the cellular detection rate (the fraction of genes that are detected with non-zero counts) as a covariate in the model. For monocle [44], the input is either TPM estimates or census counts, calculated from the TPMs as for edgeR and DESeq2 above. SCDE [51] is applied to rounded length-scaled TPMs, following the instructions provided in the package documentation, and p-values are calculated from the provided z-scores. Seurat [52] is applied using either the default “bimod” likelihood ratio test [53] (applied to the length-scaled TPMs, which are log-normalized internally), both with default settings and disabling the internal filtering steps, as well as after setting the internal expression threshold to 2 instead of the default of 0, or the “tobit” test [44] (applied to the TPMs).

Given the similarities between single-cell RNA-seq data and operational taxonomic unit (OTU) count data from 16S marker studies in metagenomics applications, we also apply metagenomeSeq [54] to the count values, fitting the zero-inflated log-normal model using the *fitFeatureModel* function from the metagenomeSeq package and testing for differences in abundance.

Finally, we include ROTS (reproducibility-optimized test statistic) [55, 56], which is a general test, originally developed for microarray data, in which a t-like test statistic is optimized for reproducibility across bootstrap resamplings. We apply ROTS to CPM and TPM values, as well as to the log-transformed CPM values calculated by the voom function in the limma package [16].

All code used for the differential expression analysis and evaluation is accessible via https://github.com/csoneson/conquer_comparison.

### Evaluation strategies

Most of the evaluations in this study are performed using real, experimental data, where no independently validated truth is available. The advantage of this approach is that no assumptions or restrictions are made regarding data distributions or specific structures of the data. However, the set of evaluation measures is more limited than in situations where the ground truth is accessible. Our first battery of evaluation approaches aim to catalog the number of genes found to be significantly differentially expressed, as well as the number and characteristics of the false positive detections from each method. For the latter evaluations, we use the null data sets, where no truly differentially expressed genes are expected and thus, all significantly differentially expressed genes are false positives. First, we investigate the fraction of genes for which no interpretable test results are returned by the applied methods (e.g., due to internal filtering or convergence failure of fitting procedures). Then, for all methods returning nominal p-values, we calculate the fraction of performed tests that give a nominal p-value below 0.05. For a well-calibrated test, this fraction should be around 5%. Next, we calculate characteristics such as the expression level (CPM), the fraction of zero counts and the expression variability (variance and coefficient of variation for CPM estimates) for all genes, and compare these characteristics between genes called differentially expressed (with an adjusted p-value/FDR threshold of 0.05) and genes not considered differentially expressed, for each of the methods. More precisely, for each characteristic and for each method detecting at least five differentially expressed genes at this threshold, we calculate a signal-to-noise statistic (*μ_S_* – *μ_NS_*)/(*σ_S_* + *σ_NS_*) where *μ_S_* (*μ_NS_*) and *σ_S_* (*σ_NS_*) represent the mean and standard deviation of the gene characteristic among the significant (nonsignificant) genes. Genes with non-interpretable test results (e.g., NA p-values) are considered non-significant in this evaluation. This approach gives insights into the inherent biases of the different methods, in the sense of the type of genes that are preferentially called significantly differentially expressed. Note that since the evaluation is done on the null data sets, the results are not confounded by the characteristics of *truly* differentially expressed genes.

The second type of evaluations focus on *robustness* of methods when applied to different subsets of the same data set. In a data set where there is a true underlying signal (i.e., truly differentially expressed genes between cell populations), ideally, this signal will be detected regardless of the set of cells that are sampled for the analysis. Thus, a high concordance between results obtained from different subsets of the cells is positive, and indicative of robust performance. For a data set without truly differentially expressed genes, however, any detections should be random, and a high similarity between results obtained from different subsets can rather indicate a bias in the differential expression calling. Thus, we first calculate a measure of concordance between the gene rankings from each pair of instances of a data set with the same number of cells per group (five such instances were generated for each group size, giving 10 pairwise comparisons). Then, we match “signal” and null instances from the same original data set and with the same number of cells per group, and calculate a t-statistic between the 10 concordance values for the signal and null instances. Thus, a large value of this statistic indicates a significant difference between the cross-instance concordance in a data set with a true underlying signal and a data set without a true signal, suggesting that the method is able to robustly detect underlying effects, and that this robustness is not due to a strong bias in the significance testing. As a measure of concordance, we use the area under the so-called *concordance curve* for the top-*K* genes ranked by significance, with *K* ∈ {100,1000}. More precisely, for each data set instance and each differential expression method, we rank the genes by statistical significance (nominal p-value or adjusted p-value/FDR estimate). Then, for each pair of data set instances with the same sample size, for *k* = 1,…, *K*, we count the number of genes that are ranked among the top *k* in both the corresponding rankings. Plotting the number of shared genes against *k* gives a curve, and the area under this curve is used as a measure of the concordance. To obtain more interpretable values, we divide the calculated area with the maximal possible value (*K*^2^/2). Thus, a normalized value of 1 indicates that the two compared rankings are identical, whereas a value of 0 indicates that the sets of top-*K* genes from the two rankings don’t share any genes. The rationale for using this type of concordance index to evaluate robustness is that it is independent of the number of genes that are actually called significant (which can vary widely across methods), and it is applicable to situations where not all compared rankings have interpretable results for the same sets of genes (e.g., due to different internal filtering criteria), which would cause a problem for e.g. overall correlation estimation. Furthermore, as opposed to a simple intersection of the top-*K* genes in the two rankings, the concordance score incorporates the actual ranking of these top-*K* genes.

A similar approach is used to evaluate similarities between methods. Briefly, for each data set instance, we rank the genes by significance using each of the differential expression methods. Then, for each pair of methods, we construct a concordance curve and calculate the area under this curve as a measure of similarity between the results from the two methods. This evaluation is only performed on the “signal” data sets.

Finally, we use the simulated data to evaluate false discovery rate (FDR) control and true positive rate (TPR, power), as well as the area under the receiver operating characteristic (ROC) curve, indicating the ability of a method to rank truly differentially expressed genes ahead of truly non-differentially expressed ones. For the pre-filtered data sets, we limit the evaluation to the genes retained after the filtering.

An interesting aspect, although not strictly related to performance, is the computational time requirement for the different methods. We investigate two aspects of this: first, the actual time required to run each method using a single core. Since this depends on the size of the data set, we normalize all times for a given data set instance so that the maximal value across all methods is 1. Thus, a “relative” computational time of 1 for a given method and a given data set instance means that this method was the slowest one for that particular instance, and a value of, e.g., 0.1 means that the time requirement was 10% of that for the slowest method. Second, we investigate how the computational time requirement scales with the number of cells. This is particularly important for scRNA-seq data, since the number of cells sequenced per study is now increasing rapidly [58]. For this, we consider all instances of all data sets (“signal” and null, as well as simulated data), and divide them into 10 equally sized bins depending on the total number of tested genes. Within each such bin, we model the required time T as a function of the number of cells per group (*N*) as

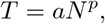

and record the estimated value of *p*.

### Software specifications

The data sets currently available in the *conquer* repository were processed with Salmon v0.6.0 [13], FastQC v0.11.6.devel and MultiQC v0.8 [24]. All analyses for the method evaluation were run in R v3.3.1 [59], with Bioconductor v3.4 [60]. Performance indices were calculated with iCOBRA v1.2.0 [61] when applicable, and results were visualized using ggplot2 v2.2.1 [62].

## Results and Discussion

### Number of differentially expressed genes

Using all instances of the seven real “signal” scRNA-seq data sets, we first compare the number of differentially expressed genes called by the different statistical methods with an adjusted p-value cutoff at 0.05. Since this number depends strongly on the number of cells per group and the strength of the signal in the data set, we represent the results as *relative* numbers, by dividing the number of differentially expressed genes detected by a given method in a given data set instance by the largest number of differentially expressed genes detected by
any method in that particular data set instance (Figure 1). From this representation, it is clear that there are large differences between the number of significant genes detected by different methods, but also that the relative ranking of methods in terms of this number varies strongly between data set instances. A breakdown by data set is shown in Supplementary Figures 4-7. In general, SeuratBimod (without the internal filtering) and monocle find a large number of significant genes, whereas SeuratBimodIsExpr2, the methods applied to census counts, ROTS, metagenomeSeq and MAST with detection rate as a covariate consistently detect much fewer genes.

**Figure 1:**
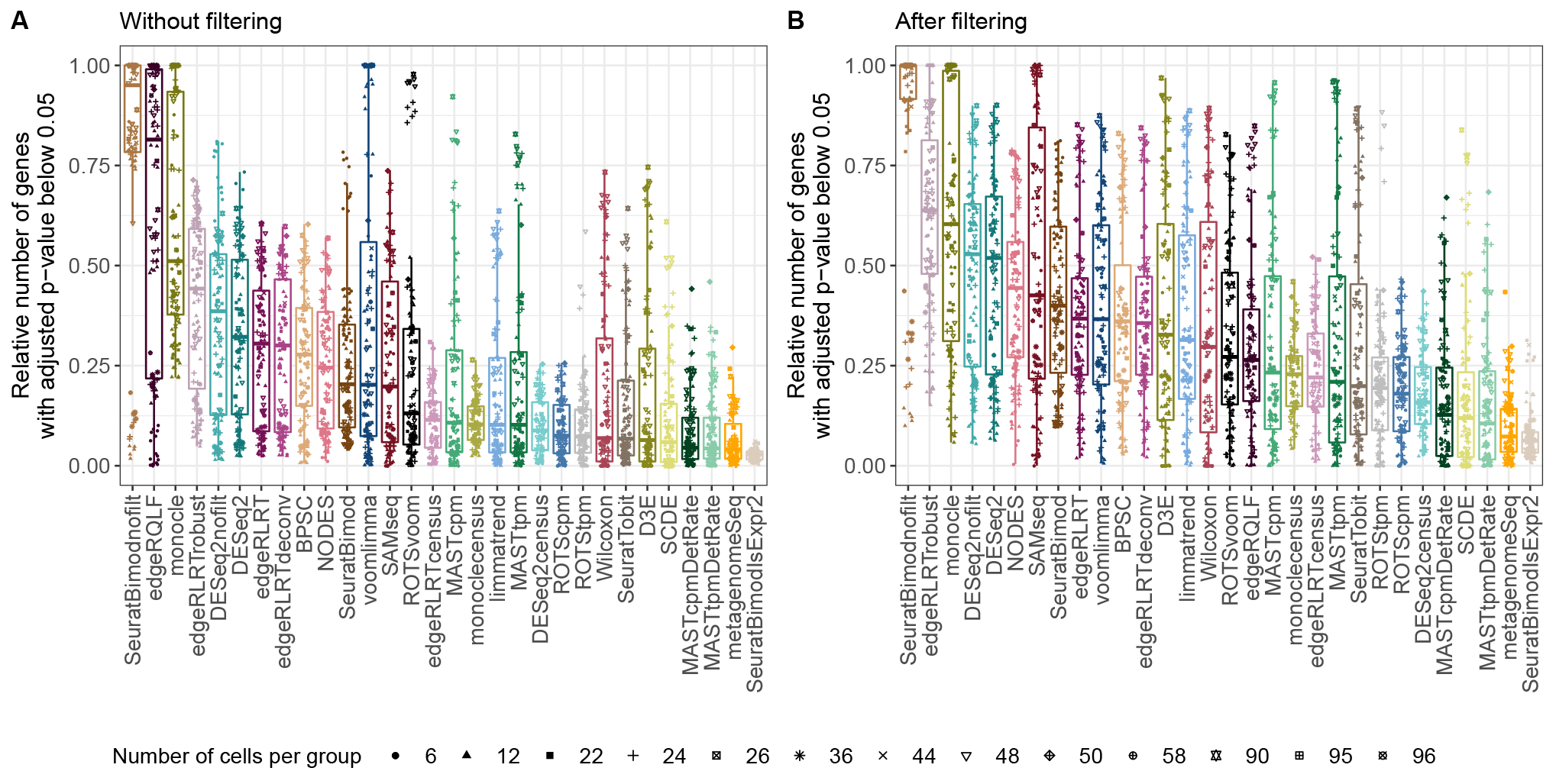
Relative number of genes detected as significantly differentially expressed (adjusted p-value below 0.05) across all instances of the seven real signal scRNA-seq data sets. For each instance, the numbers of differentially expressed genes detected by the different methods are rescaled so that the largest value is 1. The methods are ranked by the median relative number of differentially expressed genes across all instances. A. Without any prefiltering of genes, only excluding genes with zero counts across all cells. B. After filtering, retaining only genes with an estimated expression exceeding 1 TPM in more than 25% of the cells.

### Internal filtering and non-tested genes

Many of the differential expression methods, for bulk as well as single-cell analysis, implement internal filtering steps, which means that not all genes that are quantified and provided as input to the package are actually being tested for differential expression. Such filtering steps are typically performed to exclude lowly expressed genes and increase the power to detect differences in the retained genes [63, 64]. Another reason for non-reported results for a subset of the genes can be that the model fitting procedure fails to converge. Regardless of the cause, the exclusion of a subset of the genes from the reported result list can have important effects on downstream evaluations. Figure 2A illustrates the fraction of genes, for each method, that are not assigned a valid adjusted p-value or FDR estimate, aggregated across all instances of the 13 real scRNA-seq data sets (in total 188 instances with a range of sample sizes, see Supplementary Figure 8 for a breakdown by data set). While most of the methods report valid adjusted p-values for all genes, some methods exclude a large number of genes from the testing if run with default settings. For DESeq2, NODES and Seurat, this is due to explicit internal filtering of genes and/or cells, which can be disabled (corresponding to the “nofilt” methods in Figure 2A, note however that NODES failed to run when the internal filtering was disabled). Even without internal filtering, DESeq2 returns some NA p-values, corresponding to genes with low expression levels that, when the estimated counts are rounded before being input to DESeq2, have zero (rounded) counts for all cells. For metagenomeSeq, the NA p-values arise

**Figure 2:**
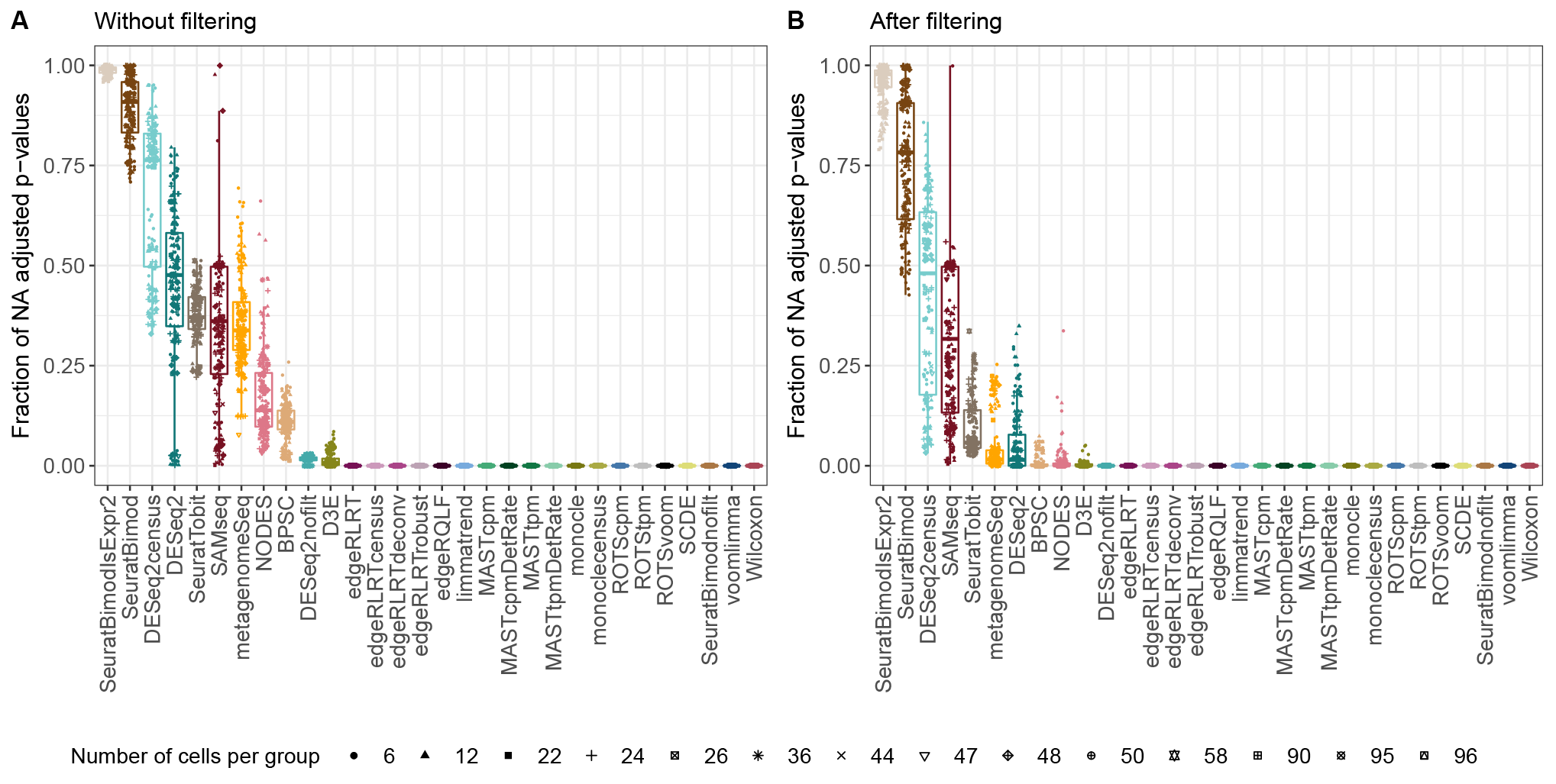
Fraction of adjusted p-values reported as NA by the different methods across 188 instances from 13 real single-cell data sets. A. Without any pre-filtering of genes, only excluding genes with zero counts across all cells. B. After filtering, retaining only genes with an estimated expression exceeding 1 TPM in more than 25% of the cells.

predominantly from genes with 0 or 1 non-zero expression values in at least one of the conditions. SAMseq only reports genes with estimated FDR strictly less than 1, and the remaining genes are excluded from the result list. For BPSC and D3E, the non-reported adjusted p-values result from convergence problems for a subset of the genes. It is worth noting that the large number of NA p-values is not a particular feature of scRNA-seq data, and similar patterns can be seen if a subset of the methods are applied to the Geuvadis (bulk) RNA-seq data set (Supplementary Figure 9).

If the data sets are filtered before the differential expression methods are applied (as described in the Methods section, retaining only genes with a TPM exceeding 1 in more than 25% of the considered cells), a large fraction of the non-reported results disappear, indicating that they mostly correspond to lowly expressed genes (Figure 2B). However, DESeq2, and in particular Seurat, still return a large number of NA test results, suggesting that their internal filtering is stricter than the fixed prefiltering criterion that we employed. For both filtered and unfiltered data, DESeq2 excludes more genes when applied to census counts than when applied to approximate read counts, likely since the former are much lower.

### Type I error control

Using the six real null data sets, where no truly differentially expressed genes are expected, we evaluate the type I error control for the methods returning nominal p-values, by recording the fraction of tested genes (with a valid p-value) that are assigned a nominal p-value below 0.05 (Figure 3A). Ideally, this fraction should be close to 5%, indicated by the horizontal line. When provided with a data set containing the full, unfiltered set of genes, many methods struggle to correctly control the type I error. The best performance is obtained by ROTS and SeuratTobit, which robustly control the type I error close to its nominal level. Several of the other methods are too liberal, with SeuratBimod and edgeR/QLF standing out with a large number of false positive findings. Setting a non-zero expression threshold in Seurat (SeuratBimodIsExpr2) improves the error control, however, as seen above (Figure 1), at the price of detecting much fewer genes as significant. Conversely, metagenomeSeq, SCDE and DESeq2 on census counts instead are too conservative. Interestingly, voom-limma mostly performs well, but for some data set instances the number of false positives is very high. A closer examination of these instances suggest that a large FPR is linked to a large set of genes with low expression, inducing a non-monotonic relationship between the mean and variance after the voom transformation (Supplementary Figure 10), affecting the observation weights and inducing a shift of the log-fold-changes of the entire set of genes. This example illustrates the practical importance of appropriate filtering as well as a visual inspection of the results. After filtering out lowly expressed genes (Figure 3B), the performance of edgeR/QLF improves dramatically, along with most other methods, while SeuratBimod still assigns low p-values to a large fraction of the tested genes. Notably, as shown in Figure 2, SeuratBimod is also among the methods returning the largest number of NA results, implying that the total number of tested genes (the denominator in the FPR calculations in Figure 3) is small. However, even when the internal filtering is disabled (SeuratBimodnofilt), the type I error is higher than for most other methods. P-value histograms confirm these observations and illustrate that without filtering, few methods return uniformly distributed p-values while after the applied filtering, results are considerably improved (Supplementary Figures 11-12). The results are largely similar for the three simulated data sets, indicating that the aspects affecting the type I error control are well captured by the simulations (Supplementary Figure 13).

**Figure 3:**
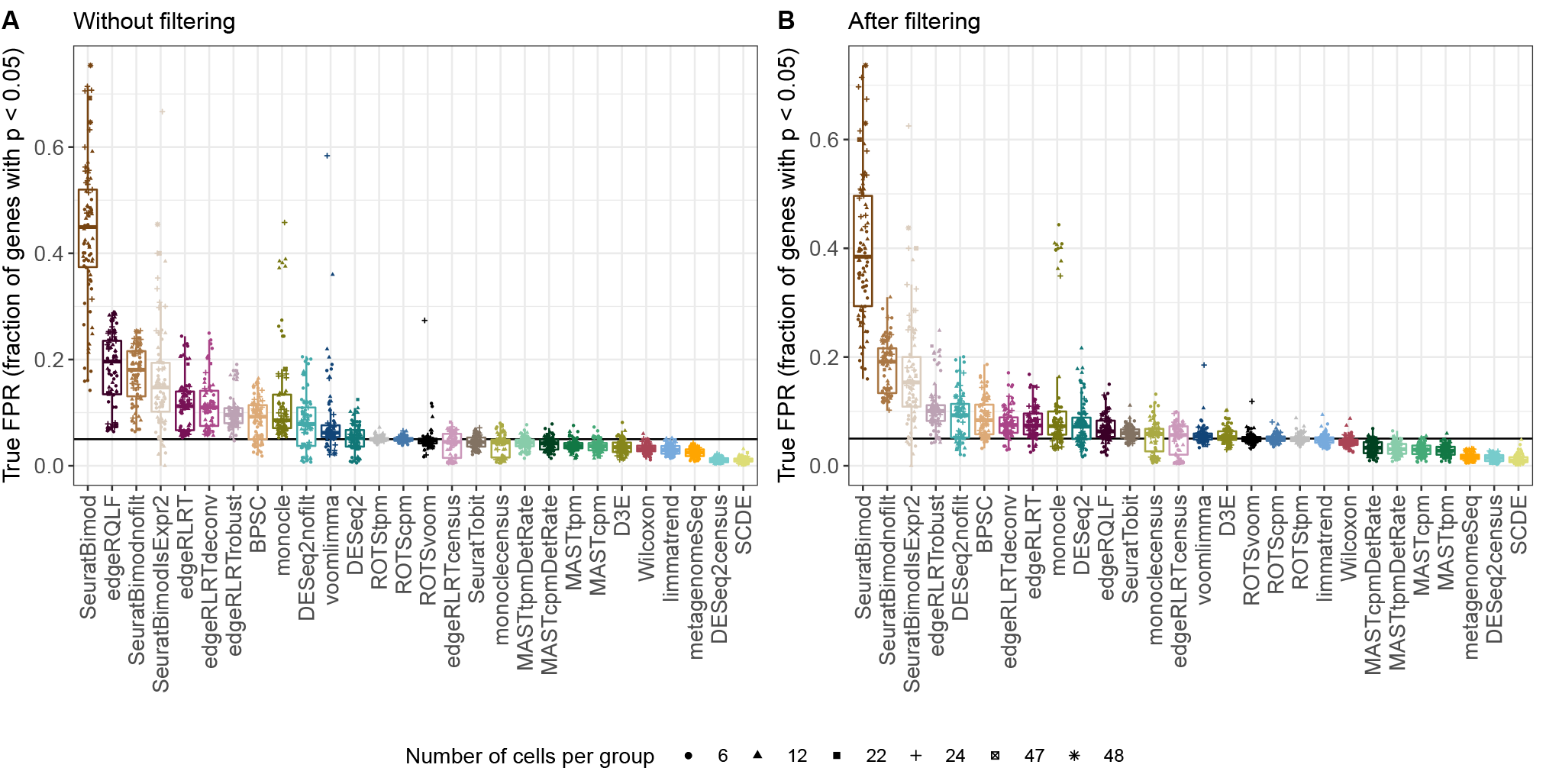
Type I error control across several instances from each of six single-cell null data sets, with a range of sample sizes. (A) Without any pre-filtering of genes. (B) After filtering, retaining only genes with an estimated expression above 1 TPM in more than 25% of the cells.

Using census counts instead of the actual estimated counts reduces the type I error for both edgeR, DESeq2 and monocle. A likely explanation for this is that the library sizes for the census count matrices are much lower than for the original count matrices, reducing the number of genes that are called significant (Figure 1).

### Characteristics of false positive genes

Knowing the biases of a particular method is important when performing differential expression analysis on real data. To investigate this aspect, we use the six unfiltered real null data sets to characterize the set of genes that are (falsely) called significant by the different methods. More precisely, for each gene in each instance of the six data sets, we estimate the average, variance and coefficient of variation of the CPM values across all cells as well as the fraction of cells in which the gene is not detected (i.e., where it has an estimated expression of zero). For each data set instance, and for each method calling at least five genes significantly differentially expressed, we then calculate a signal-to-noise statistic comparing the values of each of the four gene characteristics between the significant and non-significant (including non-tested) genes (Figure 4). The results illustrate striking differences between the types of genes detected with the different methods. The false positives detected by NODES, ROTS, SAMseq and SeuratBimod (also with non-zero expression threshold) have a low fraction of zeros, a high expression and, except for SeuratBimod with default settings, a relatively low coefficient of variation. Conversely, the false positives of edgeR/QLF, SeuratTobit, monocle, MAST and metagenomeSeq are genes with a relatively large fraction of zeros. monocle and SeuratTobit appear further biased towards genes with low expression and low variance, whereas the false positives of edgeR/QLF and MAST show higher expression and variance. The methods based on census counts preferably detect genes with high CPM values. Note that ROTSvoom, the Wilcoxon test, limma-trend and D3E did not give enough false positive findings to be included in this evaluation. Also here, the same evaluation performed on the simulated data sets shows largely similar results (Supplementary Figure 14).

**Figure 4:**
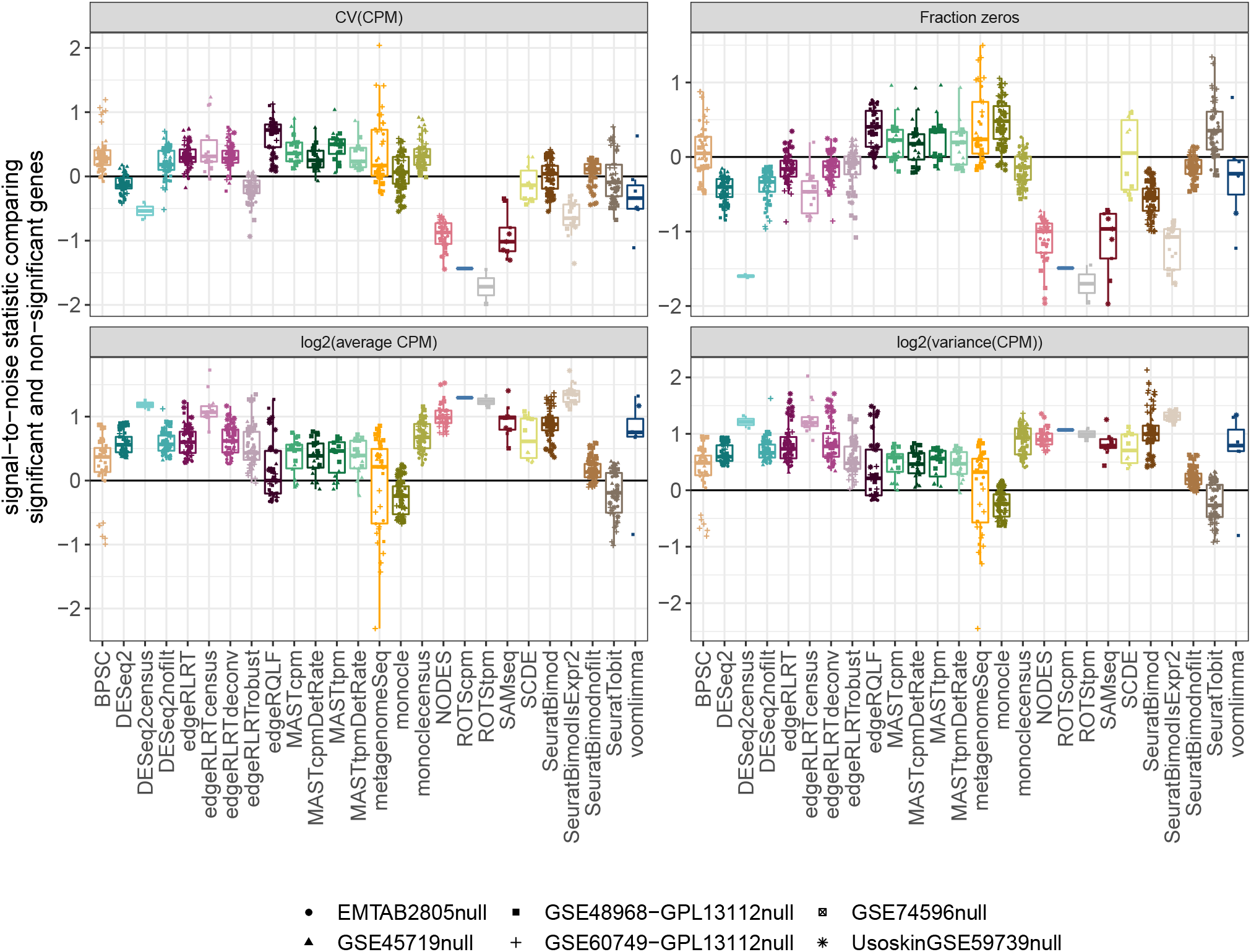
Characteristics of genes falsely called significant by each of the evaluated methods. For each instance of the six real scRNA-seq null data sets, we record characteristics of each gene (average CPM, variance and coefficient of variation of CPM, fraction zeros across all cells) and use a signal-to-noise statistic to compare each of these characteristics between genes called significant and the rest of the genes. A positive statistic indicates that the corresponding characteristic is more pronounced in the set of genes called significant than in the remaining genes. Note that ROTSvoom, D3E, limma-trend and the Wilcoxon test did not return enough false positive findings to be included in the evaluation.

### Within-method consistency

Here we investigate the notion that a good differential expression method should show high concordance between gene rankings obtained when applied to different subsets of a data set with a true underlying signal, and low concordance of rankings obtained for subsets of a null data set without a true signal. For each pair of same-sample-size instances of the six scRNA-seq data sets where both signal and null subsets are available, we generate concordance curves between the gene rankings obtained by each method. We calculate the area under each such curve, up to *K* = 100, and use this number to represent the concordance between the rankings (Figures 5A-B). Calculating a t-statistic, comparing concordances seen between subsets of the filtered “signal” data sets and the ones seen between subsets of the corresponding null data set with the same number of cells per group (Figure 5C), it is clear that all methods are generally more consistent for the signal data sets than for the null data sets, but the differencs between the methods are often small and as expected, the concordances depend strongly on the data set. Most methods achieve the best performance for the GSE60749-GPL13112 data set where the signal is very strong, and the smallest difference between signal and null data sets is seen for the EMTAB2805 data set, where the signal is much less pronounced (Supplementary Figure 1-2). Concordance values for each data set are shown in Supplementary Figure 15. Interestingly, non-parametric methods (Wilcoxon test, SAMseq) and methods based on log-like transformed data (limma-trend, ROTSvoom, voom-limma) show a slight advantage over many count-based methods in terms of the median concordance difference between signal and null data sets, which may indicate that count-based methods are more sensitive to small perturbations of the scRNA-seq data. Overall similar results are seen when using a larger value of *K* for the concordance calculations (Supplementary Figure 16).

**Figure 5:**
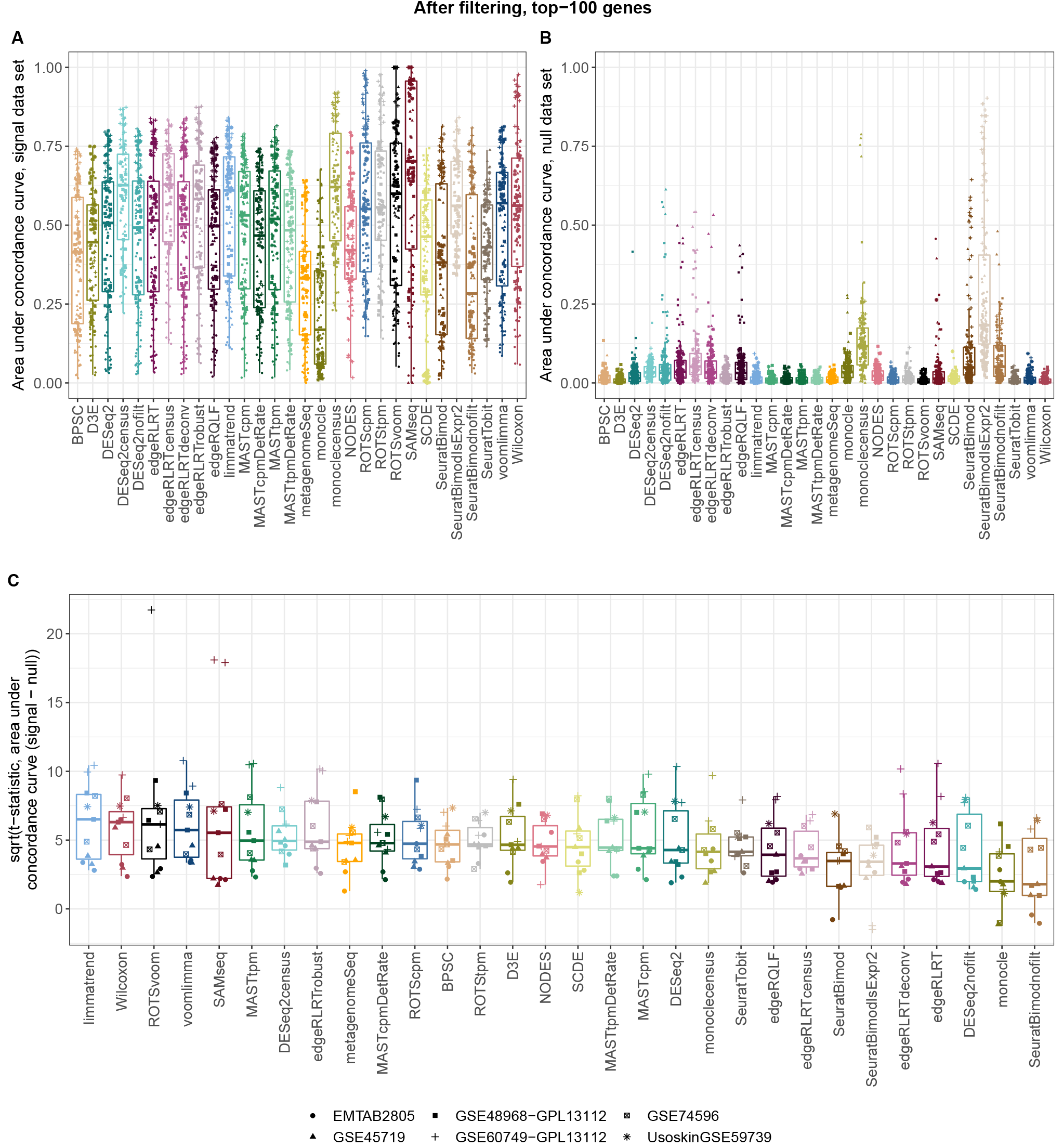
Comparison of concordance scores for the real scRNA-seq signal and null data sets. A concordance curve is constructed between the rankings of each method obtained for each pair of data set instances with the same sample size, and the concordance score is defined as the partial area under the concordance curve, until *K* = 100, divided by the maximal possible value (*K*^2^/2) to obtain relative areas between 0 and 1. A. Area under the concordance curve for the signal data sets. B. Area under the concordance curve for the null data sets. C. t-statistic comparing the areas for the signal and null data sets with the same number of cells per group. For improved visibility, the values shown are 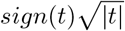.

### Between-method consistency

Using the seven scRNA-seq “signal” data sets, we quantify the concordance between gene rankings returned by different statistical methods. For each data set instance, we calculate the area under the concordance curve (AUCC) for the top-ranked 100 genes for each pair of methods. Averaging the AUCCs across all data set instances and clustering the methods based on the resulting similarity vectors (dendrogram in Figure 6) shows among other things that while the four MAST modes give overall similar rankings, the inclusion of the detection rate as a covariate has a larger effect on the rankings than changing the type of expression values from CPMs to TPMs. Moreover, the traditional count-based bulk RNA-seq methods cluster together, as do the general non-parametric methods (the Wilcoxon test and SAMseq), which are also similar to the robust count-based methods and the approaches based on log-like transformations of the data. The methods using census counts as input give similar rankings, suggesting that the change in expression unit from read counts to census counts can have a large impact on the gene rankings. Considering the distribution of individual AUCC values underlying this average across data set instances (color bars in Figure 6) illustrates that the degree of similarity between any given pair of methods can vary widely across the data set instances. For most method pairs, the degree of similarity is not strongly associated with the number of cells per group (Supplementary Figure 17) or the average silhouette width of the cells [65] (Supplementary Figure 18), and overall similar results are obtained with a larger value of *K* (Supplementary Figure 19).

**Figure 6:**
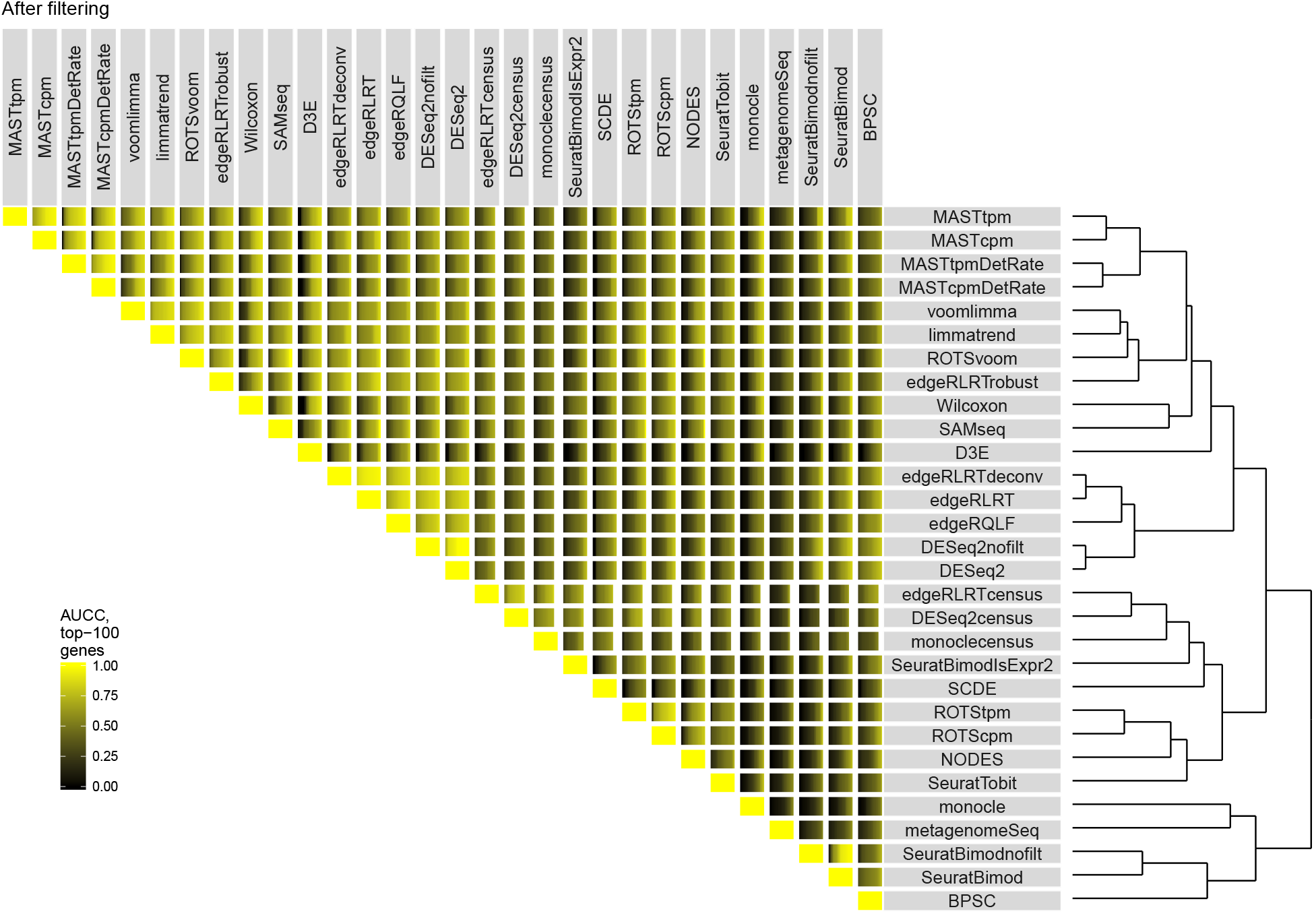
Distribution of the area under the partial concordance curve (until K = 100) for each pair of methods, across all data set instances. Dark colors indicate low concordance and yellow indicates high concordance, and each cell shows the concordance scores across all considered data set instances. The dendrogram is obtained by complete-linkage hierarchical clustering based on pairwise Euclidean distances calculated from the matrix of average AUCC values across all data set instances.

### FDR control and power

Based on the three simulated data sets, we evaluate the false discovery rate control and statistical power of the various methods. FDR control is quite poor for many methods, especially for the unfiltered data sets (Figure 7A). However, several methods, among those ROTStpm, MAST, DESeq2 applied to census counts, SeuratTobit, SeuratBimod with non-zero expression cutoff and SAMseq, robustly control the FDR close to the imposed level. SCDE, D3E, limma-trend, the Wilcoxon test, and the other variants of ROTS are somewhat conservative. The worst FDR control for the unfiltered data are obtained for monocle, SeuratBimod and edgeR/QLF. After filtering, edgeR/QLF improves dramatically (Figure 7B), now controlling the FDR close to the desired level. Also metagenomeSeq and voom-limma achieve the imposed FDR more robustly after filtering, whereas MAST and SCDE become more conservative. Comparing achieved FDRs across sample sizes shows that most methods perform closer to the optimal level for large sample sizes (Supplementary Figure 20). Non-parametric methods, such as SAMseq and the Wilcoxon test, are naturally conservative for small numbers of cells, but achieve close to the imposed FDR for larger sample sizes.

**Figure 7:**
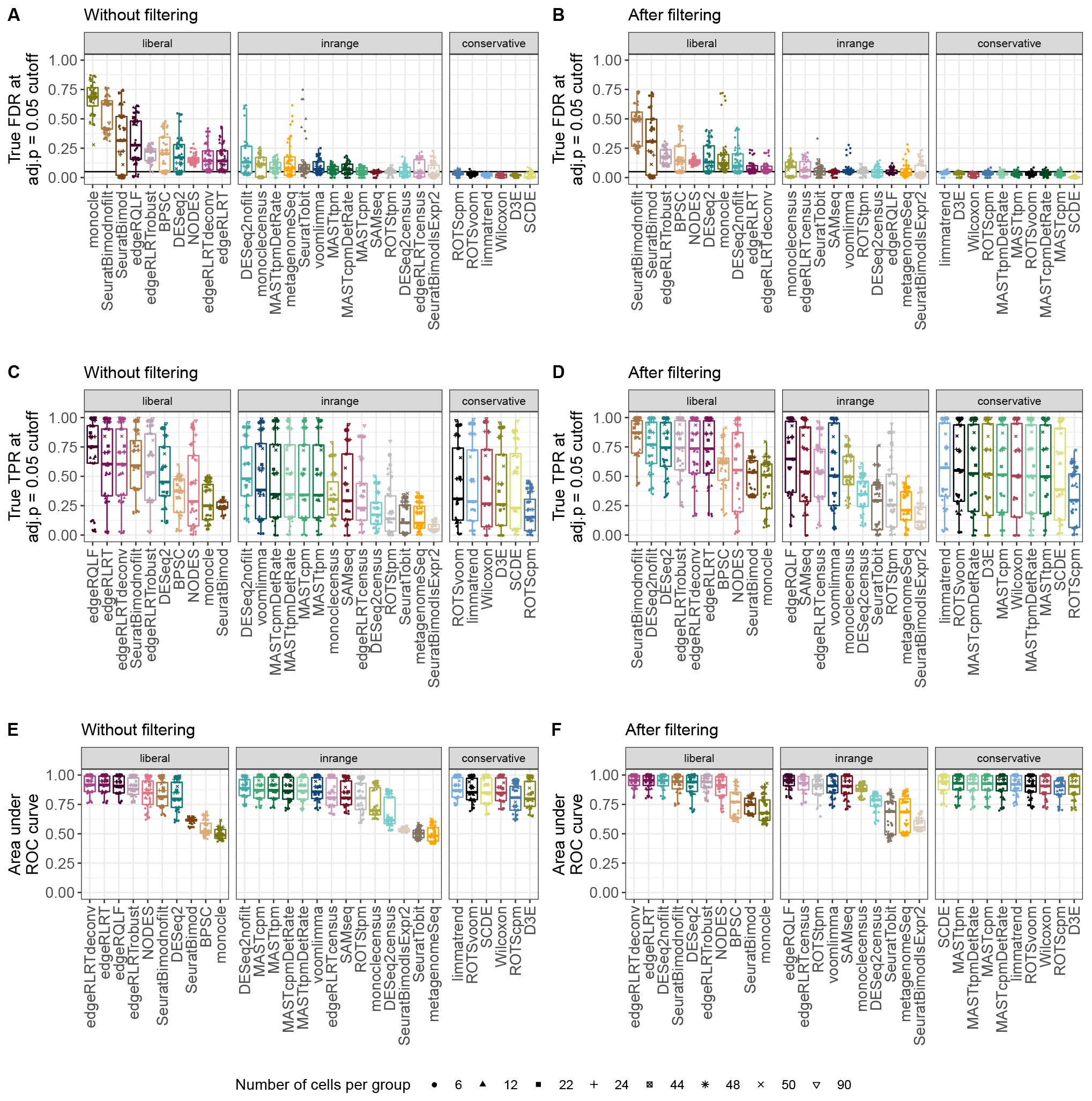
True performance, summarized across all instances of the three simulated data sets. The methods are stratified by their ability to control the FDR at the 0.05 level across the data set instances. A method where more than 75% of the observed FDRs are above 0.05 or where the median FDR is above 0.15 is considered “liberal”, whereas a method where more than 75% of the observed FDRs are below 0.05 or where the median FDR is below 0.0167 is considered “conservative”. A-B. Achieved FDR at an adjusted p-value cutoff at 0.05. C-D. Achieved TPR at an adjusted p-value cutoff at 0.05. E-F. Achieved area under the ROC curve.

As expected, the power to detect the true signal is highly dependent on the number of cells per group (Figure 7C-D), and practically all methods show increased power as the sample size grows (Supplementary Figure 21). Unsurprisingly, the largest power is obtained by some of the methods that are considered too liberal based on their FDR control. Among the methods with good, robust FDR control after filtering, edgeR/QLF, SAMseq and voom-limma achieve high power, whereas for methods like metagenomeSeq, SeuratTobit, SeuratBimodIsExpr2 and the methods applied to census counts, the FDR control comes at the price of reduced power. Also some of the more conservative methods, like limma-trend, ROTS, D3E, MAST and the Wilcoxon test, achive power comparable to less conservative methods.

The area under the ROC curve, indicating whether the methods are able to rank truly differentially expressed ahead of truly non-differentially expressed ones, shows favourable performance of edgeR (both LRT and QLF versions), followed by MAST, limma (voom and trend), SCDE, DESeq2 and SeuratBimod without filtering and the non-parametric methods (Figure 7E). Again, after prefiltering the rankings of most methods are improved (Figure 7F), and the AUROC is typically higher for data sets with larger number of cells per group (Supplementary Figure 22).

### Computational time requirement

As the number of cells included in single-cell RNA-seq data sets grows larger, the computational time requirement, the ability to parallelize calculations, and scalability become important factors in method selection. Figure 8A shows a summary of the time requirement for the different methods across all data set instances. To make values comparable across data sets with different number of cells and genes, we scale the times for each data set instance relative to the slowest method for that particular instance to get relative time requirements. All methods were run on a single core. Three dedicated single-cell methods, namely BPSC, D3E and SCDE, stand out as being the slowest for most of the data sets, while the bulk methods (edgeR, DESeq2 and especially the limma variants) are generally considerably faster. However, many of the methods are easy to parallelize, which can speed up the calculations. In particular, DESeq2, BPSC, MAST, SCDE and monocle all feature explicit arguments to take advantage of parallelization. In addition, methods that are performing gene-wise tests without information sharing between genes, such as the Wilcoxon test and D3E, can be manually run in parallel after splitting the data into multiple chunks. The computational time required by Seurat depends strongly on whether its built-in filtering or a non-zero expression threshold is used, in which case a large number of the genes are excluded from testing.

**Figure 8:**
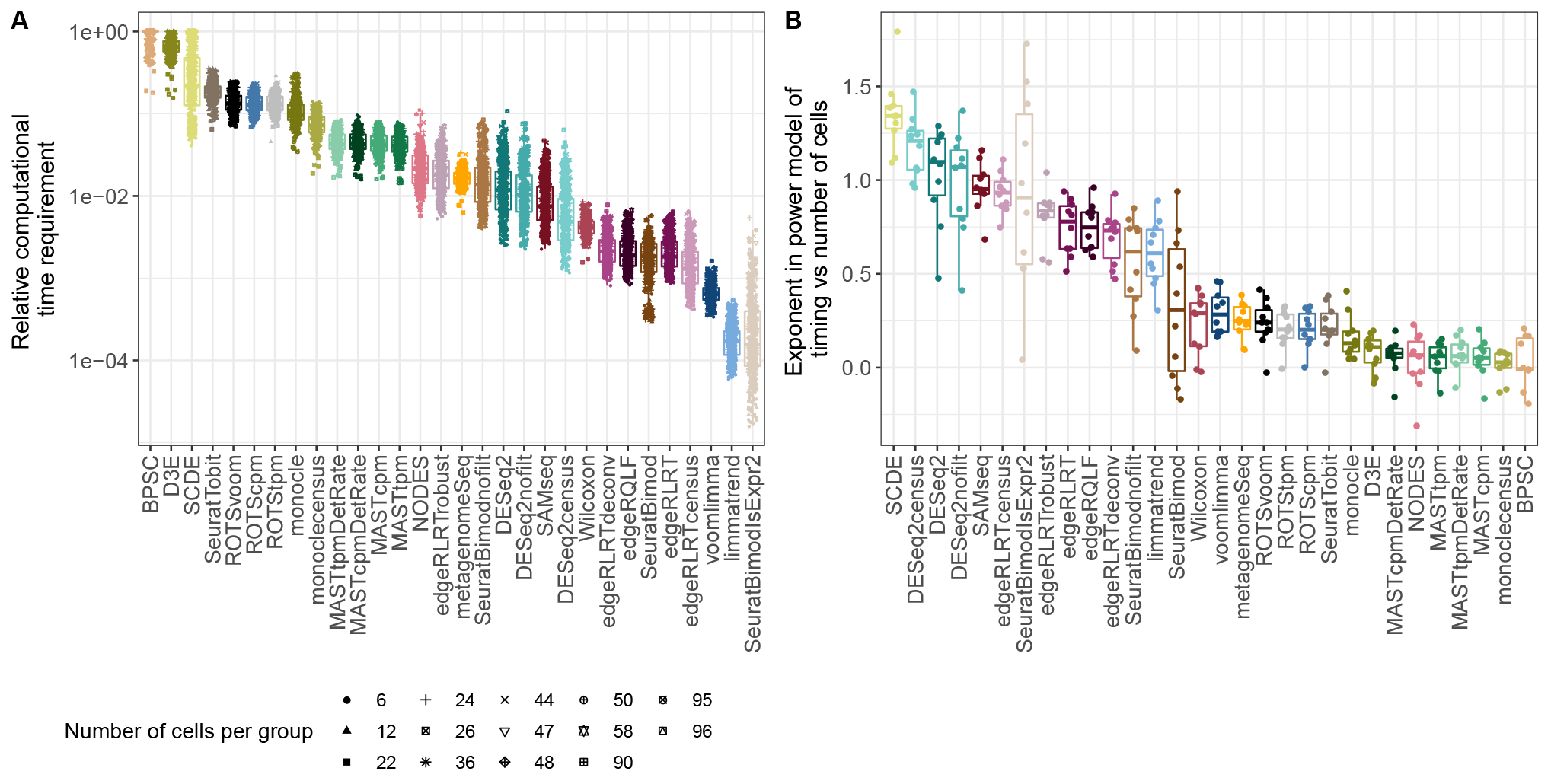
A. Relative computational time requirement for the different methods across all scRNA-seq data sets. B. Exponent in a power model relating computational time requirement to the number of cells, for data sets instances with similar number of genes.

In addition to the actual time required for the calculations, we evaluate how the methods scale with increasing number of cells. While the number of genes will remain roughly the same, the number of sequenced cells is likely to increase in the future and it is important that computational methods can accommodate this. In this respect, there is a relatively clear distinction between methods developed for bulk RNA-seq analysis and those developed explicitly for single-cell RNA-seq data. In particular, most single-cell methods (with the exception of SCDE) scale very well with increasing number of cells, while the computational time required for the bulk RNA-seq methods is more dependent on the sample size (Figure 8B). Supplementary Figures 23-26 summarize the actual computational time for each method across all data set instances, and Supplementary Figure 27 illustrates the fitted power models of the computational time as a function of the number of cells per group for each collection of data set instances with similar number of genes.

### Other aspects

While the evaluations in this study are centered on the simplest experimental situation, comparing two groups of cells, many real studies require the formulation of a more complex experimental design. Currently, not all evaluated methods can accommodate such situations. Specifically, the Wilcoxon test, NODES, SCDE, Seurat, ROTS and D3E are limited to two-group comparisons, and SAMseq can only accommodate a limited number of experimental designs. The remaining methods implement statistical frameworks that can accommodate more complex (fixed effect) experimental designs, including adjustments for known batch effects and other covariates.

## Conclusions

We have presented *conquer*, a repository of consistently processed public single-cell RNA-seq data sets, and used a subset of the included data sets for an extensive evaluation and comparison of methods for differential expression analysis. The fact that *conquer* provides gene expression estimates in multiple units allowed us to compare methods requiring different types of input values, and also to investigate the effect of using different input values for the same statistical method. We have shown that prefiltering of genes is essential to obtain good, robust performance for several of the evaluated methods, most notably edgeR/QLF, which tends to call lowly expressed genes with many zeros significant if these are present in the data but otherwise performs well, and voom-limma, which also performs more robustly after filtering out lowly expressed genes. We noted a large variability among the number of genes called differentially expressed with the different methods, as well as in the ability to control the type I error rate and the false discovery rate. After appropriate filtering, a subset of the methods, most notably SAMseq, limma (voom and trend), D3E, ROTSvoom and the Wilcoxon test, but also edgeR/QLF and MAST, managed to control the FDR and FPR close to the imposed level while achieving a high power while for most other methods, appropriate error control was associated with a lack of power.

We also showed that the differential expression methods are biased in different ways in terms of the types of genes they preferentially detect as significantly differentially expressed, which can have important implications in practical applications. The results presented here can be compared to the genes detected with a given method in a real data set, to indicate possible over- or under-representation of genes with specific characteristics. In agreement with previous evaluations, methods originally developed for bulk RNA-seq analysis did not perform significantly worse than methods specifically developed for scRNA-seq data, but sometimes showed a stronger dependency on the data being pre-filtered. The non-parametric methods and methods using log-like transformations of the count data showed a slight advantage in terms of robustness across data subsets.

Throughout this study, we have used four different abundance units to represent the expression values. The TPM and approximate count (length-scaled TPM) values were directly obtained from the Salmon output, whereas CPM values were calculated from the counts using edgeR, and census counts were estimated from the TPMs using monocle. For the latter, we used the default settings of the *relative2abs* function, and applied it directly to the gene-level TPM estimates, resulting in a relatively low total number of transcript counts. It is possible that modifications of these settings, optimized for the library preparation parameters for each individual data set, would lead to larger absolute count values, and thus potentially a higher statistical power.

The number of cells in each considered cell type ranged between 6 and 96 in our data sets. While this is a relatively small number compared to the thousands of cells that can be sequenced in an actual experiment, differential expression is typically performed between sets of homogeneous cells (e.g., from a given, well-defined cell type), and these collections are likely to be much smaller. Thus, we believe that the range of sample sizes considered in our comparisons are relevant for real applications, and that it is important to know how the methods perform under these circumstances. It is possible that some methods may improve their performance when applied to much larger collections of cells. For example, SeuratBimod shows a steep decrease in the achieved FDR as the number of cells increase (Supplementary Figure 20B).

## Data availability

*conquer* is available as a Shiny application at http://imlspenticton.uzh.ch:3838/conquer/. All code used to process the data sets for *conquer* can be accessed via GitHub: https://github.com/markrobinsonuzh/conquer. The code used to perform the evaluation of the differential expression analysis methods is also available from GitHub: https://github.com/csoneson/conquer_comparison.

## Authors’ contributions

CS and MDR designed analyses and wrote the manuscript. CS performed analyses. Both authors have read and approved the final manuscript.

## Acknowledgements

We would like to acknowledge Michael I Love and Valentine Svensson for helpful online instructions regarding automated downloading of raw data from ENA (https://gist.github.com/mikelove/f539631f9e187a8931d34779436a1c01, http://www.nxn.se/valent/streaming-rna-seq-data-from-ena).

